# Proteome analysis of human visceral and subcutaneous adipocytes identifies depot-specific heterogeneity at metabolic control points

**DOI:** 10.1101/2020.04.13.039057

**Authors:** Arthe Raajendiran, Christoph Krisp, David P. De Souza, Geraldine Ooi, Paul R. Burton, Renea A. Taylor, Mark P. Molloy, Matthew J. Watt

**Affiliations:** Department of Physiology, The University of Melbourne, Melbourne, Victoria, 3010, Australia; Monash Biomedicine Discovery Institute, Department of Physiology, Monash University, Clayton, Victoria, 3800, Australia; Monash Biomedicine Discovery Institute, Metabolism, Diabetes and Obesity Program; Australian Proteome Analysis Facility, Macquarie University, NSW 2109, Australia; Metabolomics Australia, Bio21 Institute of Molecular Science and Biotechnology, University of Melbourne, Parkville 3010, Australia; Centre for Obesity Research and Education, Faculty of Medicine, Nursing and Health Sciences, Monash University, Melbourne, Victoria, 3004, Australia; Cancer Research Division, Peter MacCallum Cancer Centre, University of Melbourne, Melbourne, Victoria, 3000, Australia

## Abstract

Adipose tissue is a primary regulator of energy balance and metabolism. The distribution of adipose tissue depots is of clinical interest because the accumulation of upper-body subcutaneous (ASAT) and visceral adipose tissue (VAT) is associated with cardiometabolic diseases, whereas lower-body gluteal-femoral adipose tissue (GFAT) appears to be protective. There is heterogeneity in morphology and metabolism of adipocytes obtained from different regions of the body, but detailed knowledge of the constituent proteins in each depot is lacking. Here, we determined the human adipocyte proteome from ASAT, VAT and GFAT using high-resolution SWATH mass spectrometry proteomics. We quantified 4220 proteins in adipocytes, and 2329 proteins were expressed in all three adipose depots. Comparative analysis revealed significant differences between adipocytes from different regions (6 and 8% when comparing VAT *vs.* ASAT and GFAT, 3% when comparing ASAT *vs.* GFAT), with marked differences in proteins that regulate metabolic functions. The VAT adipocyte proteome was overrepresented with proteins of glycolysis, lipogenesis, oxidative stress and mitochondrial dysfunction. The GFAT adipocyte proteome predicted activation of PPARα, fatty acid and BCAA oxidation, enhanced TCA cycle flux and oxidative phosphorylation, which was supported by metabolomic data obtained from adipocytes from the same patient donors. Together, this proteomic analysis provides an important resource and novel insights that enhance the understanding of metabolic heterogeneity in the regional adipocytes of humans.

## Introduction

White adipocytes play a crucial role in maintaining systemic energy homeostasis by storing and releasing fatty acids in coordination with feeding and fasting cycles, respectively (Rosen and Spiegelman, 2006; Scherer, 2016). While these functions of adipose tissue are conserved across the body’s various storage sites, the capacities of adipocytes for fatty acid storage and release vary according to the anatomical location of adipose tissue depots. For example, lipolysis rates are increased in adipocytes located in subcutaneous compared with visceral regions, and within the subcutaneous adipose tissue depots, lipolysis is greater in adipocytes derived from upper-body abdominal (ASAT) compared with the lower body glutealfemoral (GFAT) adipose tissue (Demerath et al., 2007; Jensen, 2008; Rajjo et al., 2014; Salans et al., 1968; Shadid et al., 2007; Tchernof et al., 2006). There is also evidence for regional differences in the adipocytes capacity to sequester fatty acids from chylomicrons (meal-derived), very-low density lipoproteins (VLDL, secreted by the liver) and from circulating free fatty acids (FFA) (Koutsari et al., 2008; McQuaid et al., 2010; Votruba et al., 2007). It has been postulated that these alterations in lipid metabolism (McQuaid et al., 2011; Spalding et al., 2017) may partly explain the apparent contribution of visceral (VAT) (Karlsson et al., 2019) and abdominal subcutaneous (ASAT) adipose tissue depots to the pathogenesis of insulin resistance and other cardiometabolic diseases, and the possible protective role of glutealfemoral adipose tissue (GFAT) in disease pathogenesis (Lee et al., 2013; McQuaid et al., 2011).

Adipose tissues play a significant role in the regulation of glucose (Horowitz et al., 2001; Marin et al., 1987) and branched-chain amino acid (BCAA) metabolism (Felig et al., 1969; Newgard et al., 2009) (Herman et al., 2010; Rosenthal et al., 1974; Suryawan et al., 1998), and there is a growing appreciation that adipose tissues participate in a variety of other metabolic functions that impact whole body energy homeostasis and produce metabolic signals that participate in inter-tissue cross-talk (Funcke and Scherer, 2019). However, the contribution of adipocytes from different anatomical locations to these processes remains incompletely understood. Understanding how and why regional adipocytes within the same individual orchestrate the variable uptake and processing of these, and other, metabolites may be useful for predicting the overall metabolic health of individuals.

The molecular regulation within adipocytes directs the production and degradation of proteins that invariably control cellular functions (Ali et al., 2011). Others have profiled the whole transcriptome of human adipose tissues obtained from visceral and subcutaneous locations (Bradford et al., 2019; Gerhard et al., 2014; Linder et al., 2004), which indicated that the genes involved in cellular and embryonic development, connective tissues disorders, cell morphology, lipid droplet formation and mobilization and angiogenesis-related pathways were differentially regulated between these adipose sites. Metabolism is controlled by the abundance and post-translational modifications of proteins (Adachi et al., 2007). Cross-sectional analysis of the proteomes from whole VAT and ASAT (Fang et al., 2015; Insenser et al., 2012; Perez-Perez et al., 2009) indicate that proteins associated with glucose and lipid metabolism, lipid transport, response to stress and inflammation and nuclear receptor activation pathways were differentially regulated, while a recent study (Vogel et al., 2019) reported few differences between the ASAT and GFAT proteomes. However, adipose tissue is composed of many cell types including mature adipocytes, adipocyte precursors (*i.e.*, pre-adipocytes), immune cells, endothelial cells and fibroblasts (Ouchi et al., 2011), hence analyzing the whole adipose tissue proteome does not accurately represent the proteome of adipocytes. In this regard, recent studies have profiled the proteome of freshly isolated mature adipocytes obtained from ASAT of lean and obese individuals and detected ~1,500 proteins (Xie et al., 2010; Xie et al., 2016), however, comparisons between regional adipose tissue depots were not assessed. Together, these studies suggest that the metabolic phenotype of the adipose tissue is heavily influenced by the adipocyte proteome and assessing the proteome of regional adipocytes will facilitate the understanding of the depot-specific contribution to metabolic health and disease.

In the present study, we have used SWATH mass spectrometry to assess the proteome of adipocytes isolated from patient-matched upper (VAT and ASAT) and lower-body (GFAT) adipose tissues. We also performed metabolomic analysis on adipocytes derived from the same donors and integrated these data with the proteomic data to gain insights into the control of metabolism in regional adipocytes.

## Methods

### Adipose tissue collection and adipocyte isolation

Ethics approval was obtained from the Alfred Hospital Ethics Committee in accordance with the National Statement on Ethical Conduct in Human Research (2007) (Alfred Ethics Approval No. 548/14). All participants provided informed consent to participate in this study. Adipose tissue biopsies from VAT, ASAT and GFAT were obtained from five patients (BMI 47.3 ± 1.4 kg/m^2^, mean ± SEM) undergoing bariatric procedures as previously mentioned (Raajendiran et al., 2019). The adipose tissues (4-5 g) were processed separately as depicted in Figure 1A and the mature adipocytes were isolated as described previously (Raajendiran et al., 2019). The tissues were rinsed twice in phosphate buffered saline (PBS), blot dried for 5 seconds and immediately placed in Roswell Park Memorial Institute media - 1640 (RPMI-1640, ThermoFisher Scientific Australia) containing 3% (w:v) bovine serum albumin (BSA) (Sigma Aldrich, Australia), 1 mg/ml Type 2 Collagenase (Sigma Aldrich, Australia) at a ratio of 1:2 (g:ml). The tissues were minced into 5-10 mg pieces using scissors, transferred to a 37°C shaking water bath in 50 ml plastic tubes and incubated with agitation for 45 minutes. The digests were strained into separate 50 ml plastic tubes using 200 μm nylon filters (Corning, Australia) to remove any undigested tissue and the tubes were spun at 1000 x *g* for 10 minutes at room temperature (RT). The floating mature adipocytes from each tube were collected in separate 2 ml tubes (half filled) using plastic Pasteur pipettes. If required, additional 2 ml tubes were used to collect more cells. The cells were then washed with 1 ml of PBS three times at 100 x g for 3 min each at RT by displacing / aspirating PBS using a 19 G syringe, each time. The resulting cells were snap frozen in liquid nitrogen and stored in −80°C until analysis.

**Figure 1.**
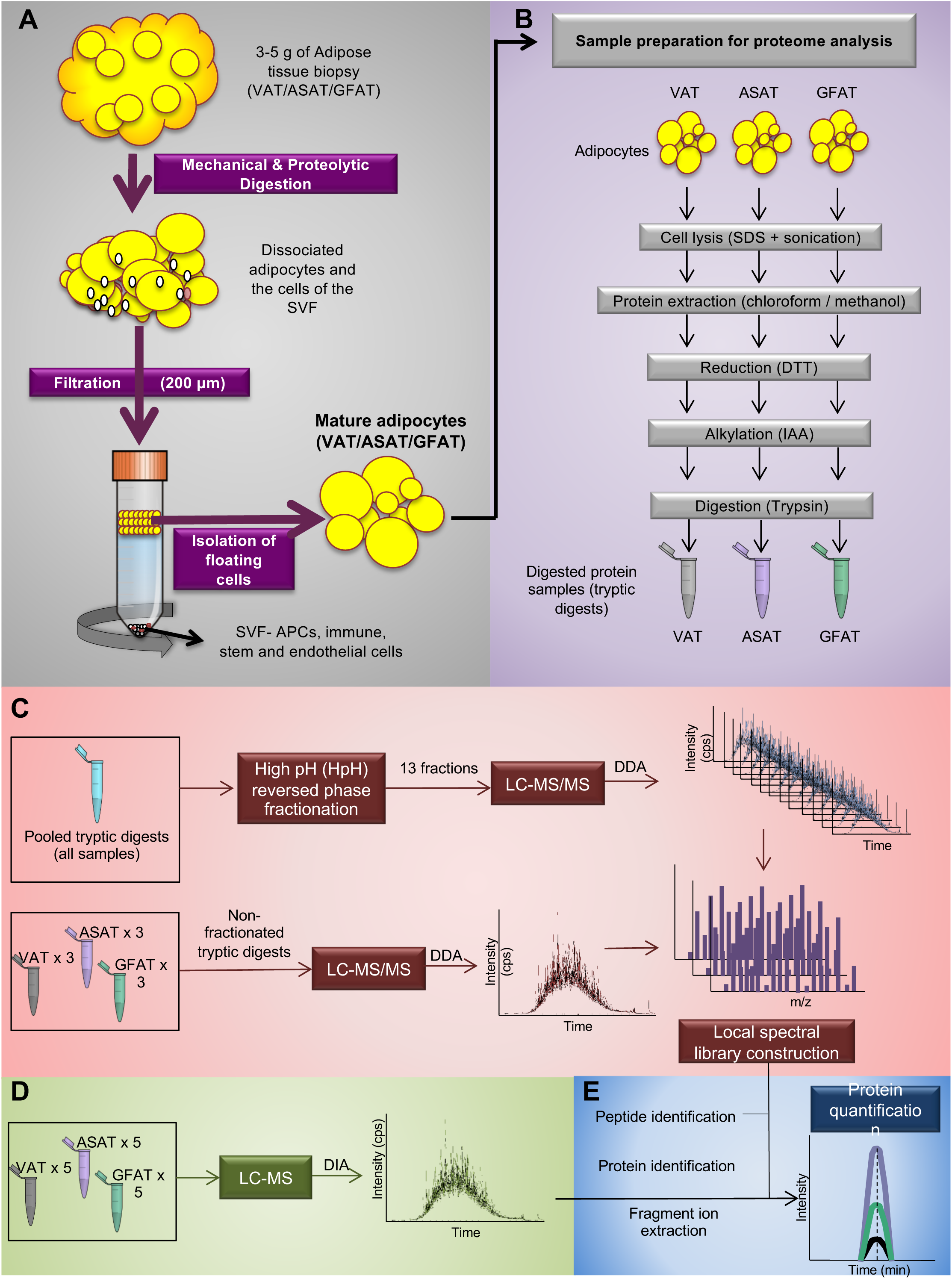
Overview of adipocyte isolation and SWATH analysis. **A.** VAT, ASAT and GFAT was obtained from patients and was digested to separate the cell populations. The cell suspension was passed through a 200 µm nylon filter to separate the mature adipocytes from the stromal vascular fraction (SVF) cells and the isolated adipocytes were snap-frozen. **B.** The adipocyte samples for mass spectrometry were processed as described in grey boxes to obtain the tryptic digests. **C.** For creating a local spectral library, an aliquot of tryptic digests of all samples (all three depots from all five subjects) was pooled and separated into thirteen fractions. Also, non-fractionated tryptic digests from three subjects per tissue type were aliquoted separately. Both fractionated and non-fractionated samples were scanned in an LC-MS/MS system by following the data-dependent acquisition (DDA) approach. The spectra were then pooled to construct a local adipocyte library. **D.** Tryptic digests from each tissue type in all subjects were then scanned separately in an LC-MS system using a data-independent acquisition (DIA) approach. **E.** The local spectral library was searched against the SWISS-prot database using ProteinPilot software to identify individual peptides and map the corresponding proteins. Fragment ions obtained from the DIA spectra of individual samples were then uploaded in the ProteinPilot software for protein quantification.

### Sample preparation for mass spectrometry

The frozen adipocytes were sent to A/Prof Mark Molloy (Australian Proteome Analysis Facility, Macquarie University, Sydney, Australia) for MS analysis. The protein digests were prepared from the frozen samples as outlined in Figure 1B. Briefly, one aliquot of adipocytes from each adipose tissue type and patient was lysed with 500 µl of 1% sodium deoxycholate in 100 mM triethylammonium bicarbonate using a probe sonicator (6 pulses). Lysed samples were heated at 95 °C for 5 min and divided into 200 µl aliquots. Proteins were extracted using chloroform / methanol (4:1). Protein pellets from each aliquot per sample were pooled and the total protein content were measured using BCA protein Assay kit (ThermoFisher Scientific, Australia). The protein disulphide bonds were reduced with 10 mM dithiothreitol for 30 min at 60 °C and the cysteine residues were alkylated with 20 mM iodoacetamide for 30 min at 37 °C in the dark. Fifty µg of each sample was digested with trypsin (20:1 protein to enzyme) overnight at 37 °C. After acidification of the samples to quench digestion and precipitation of sodium deoxycholate, samples were dried in a vacuum concentrator and resuspended in 2% acetonitrile and 0.1% formic acid to generate 1 µg/µl stocks (based on BCA assay) which were then subjected for LC-MS/MS and LC-SWATH-MS characterization (Figure 1C-E).

A pool of all tryptic digests derived from VAT, ASAT and GFAT of all the patients (100 µg) was generated and fractionated by basic reversed phase chromatography (HpH RP) prior to LC-MS/MS to decrease the complexity and enhance the peptide and protein identification (Yang et al., 2012). A C18 solid phase extraction was performed and the purified peptides were dried. The peptides were resuspended in Solvent A (5 mM ammonia solution, pH 10.5) and loaded onto a C18 RP column (300 Extend-C18 column, 2.1 mm x 150 mm, 3.5 µm, 300Å column, Agilent). The column was washed in a solvent gradient system involving a mixture of 97% Solvent A and 3% Solvent B (5 mM ammonia solution with 90% acetonitrile, pH 10.5) for 10 minutes. The concentration of Solvent B was subsequently increased to 30% over 55 minutes, and gradually increased to 70% within 10 minutes and finally to 90% within 5 minutes at a flow rate of 300 μl/min. The eluent was collected at two-minute intervals for the first 20 min of the gradient and at one-minute intervals for the rest of the gradient. Collected fractions were pooled into 13 fractions (3 fractions from 0 – 22 min and 10 fractions from 22 – 82 min pooling collections taken every 10 min into one fraction), dried and resuspended in 2% acetonitrile and 0.1% formic acid to give a ~0.1 μg/μl stock.

### LC-MS

LC-MS acquisition was performed as published by Martínez-Aguilar *et al* (Martinez-Aguilar et al., 2016) with minor modifications. Nine non-fractionated tryptic digests (3 x VAT, 3 x ASAT and 3 x GFAT - from different subjects) and thirteen HpH RP fractions were analyzed data dependent acquisition (DDA) on a 6600 TripleToF with an Eksigent nanoLC 415 system with cHiPLC system (SCIEX). Ten μl (0.1 μg/μl) of each sample were loaded onto a ChromXP C18 peptide trap (15 cm x 200 μm, 3 μm particle size, 120 Å, SCIEX, USA). The peptides were separated with a 60 min gradient of 5 – 40% acetonitrile. The LC eluents were subjected to positive ion nanoflow electrospray MS analysis in DDA mode to obtain a reference spectral ion library to be used for information extraction from the SWATH-MS. A time of flight (TOF)-MS survey scan was acquired (m/z 350-1500, 255 ms accumulation time). The 20 most intense multiple charged precursor ions (2^+^ - 4^+^, counts > 200 counts per second) in the survey scan were sequentially subjected to MS/MS analysis. MS/MS spectra were accumulated for 100 ms (m/z 100-1800) with rolling collision energy. SWATH or data independent acquisition (DIA) of the fifteen-tryptic digests (5 x VAT, 5 x ASAT and 5 x GFAT) was performed under the same LC conditions as DDA. A 100 variable window method was generated with window widths selected based on precursor densities in the range from 400 to 1250 m/z using the SCIEX variable window calculator tool. MS/MS of the selected windows were acquired using 30 ms accumulation times with 3.2 s duty circles.

### Protein identification

The DDA MS files were processed using ProteinPilot software (version 5.0, SCIEX, USA) with the paragon search algorithm (Shilov et al., 2007) and the MS/MS spectra were searched against the Human SwissProt database (released Feb 2016) to generate a spectral library. The ProteinPilot search results were imported into PeakView software (version 2.1) equipped with SWATH acquisition MicroApp (version 2.0) to perform targeted information extraction. Peptides shared between proteins were excluded. Information extraction was performed using up to six fragment ions per peptide, a maximum of 100 peptides per protein with a peptide confidence > 99% were allowed, the mass tolerance was set to 75 ppm, the extraction window 5 min and a false discovery rate (FDR) ≤ 1%.

### Metabolome analysis

A known volume (~50 mg) of frozen adipocytes isolated from patient-matched VAT, ASAT and GFAT adipocytes was placed in separate 1.5 ml Eppendorf tubes and subjected to cold extraction (Chloroform: Methanol: Water at a ratio of 1:3:1 v/v/v). Briefly, 100 µl of chloroform was added to the tubes and mixed gently. Then, 400 µl of methanol:water (3:1 v/v) containing 1.25µM ^13^C_6_-Sorbitol and 12.5 µM ^13^C_5_,^15^N-Valine standards was added to the tubes and spun at a maximum speed for 5 minutes at 4°C to pellet precipitated protein and cell debris. The supernatant from the tubes was then transferred into micro-vial inserts and evaporated to dryness *in vacuo* using a Christ RVC 2-33 speed vacuum concentrator.

Dried samples for targeted analysis were derivatized online using the Shimadzu AOC6000 autosampler robot. Derivatization was achieved by adding 25 µL of Methoxyamine Hydrochloride (30 mg/mL in Pyridine) followed by shaking at 37°C for 2h. Samples were then derivatized with 25 µL of *N*,*O*-*bis* (Trimethylsilyl)trifluoroacetamide with Trimethylchlorosilane (BSTFA with 1% TMCS, Thermo Scientific) for 1h at 37°C. The sample was left for 1 h before 1 µL was injected onto the GC column using a hot needle technique. Split-less injections were performed for each sample.

The GC-MS system used comprised of an AOC6000 autosampler, a 2010 plus Shimadzu gas chromatograph and a TQ8040 quadrupole mass spectrometer (Shimadzu, Japan). The mass spectrometer was tuned according to the manufacturer’s recommendations using tris-(perfluorobutyl)-amine (CF43). GC-MS was performed on a 30m Agilent DB-5 column with 1µm film thickness and 0.25mm internal diameter column. The injection temperature (Inlet) was set at 280°C, the MS transfer line at 280°C and the ion source adjusted to 200°C. Helium was used as the carrier gas at a flow rate of 1 mL/min and Argon gas was used as the collision cell gas to generate the MRM product ion. The analysis of TMS samples was performed under the following temperature program; start at injection 100°C, a hold for 4 minutes, followed by a 10°C min^-1^ oven temperature ramp to 320°C following final hold off for 11 minutes. Approximately 520 quantifying MRM targets were collected using the Shimadzu Smart Metabolite Database (along with a qualifier MRM for each target), which covers approximately 350 endogenous metabolites and multiple ^13^C-labelled internal standards. Both chromatograms and MRMs were evaluated using the Shimadzu GCMS browser and LabSolutions Insight software. Resulting area responses for detected metabolites were exported as an area data matrix for downstream statistical analysis.

### Experimental Design and Statistical Rationale

We compared the proteomes of subject matched upper-body (VAT and ASAT) and lower-body (GFAT) adipocytes obtained from five different individuals to understand their functional differences. Individual protein peak areas (sum of areas of all peptides with FDR ≤ 1% in one of the biological replicates) obtained as mentioned in the methods section were loaded in Perseus software (version 1.5), log2 transformed and normalized by the median protein area per sample. The relative quantitation of proteins was performed between two groups using two-tailed student’s t-test. Proteins with p-values < 0.05 and with a fold-change (FC) > 1.5 or < 0.6 were considered as differentially expressed. For data visualization purposes in the figures, the log_2_-peak area of the selected ASAT and GFAT proteins per subject were normalized to their subject-matched log_2_-peak area of the VAT protein.

Likewise, the metabolite peak areas per ASAT and GFAT adipocytes per patient were normalised to the paired VAT adipocytes. The relative quantitation of normalized metabolites was performed between two groups using two-tailed student’s t-test. Metabolites with p-values < 0.05 were considered as differentially abundant.

### Bioinformatics analyses

The list of identified proteins from adipocytes derived from VAT, ASAT and GFAT were subjected to enrichment analyses using the online Gene Ontology (GO) database (www.geneontology.org) to determine their subcellular localization. Volcano plots were generated using GraphPad prism software and heat maps and principle component analysis (PCA) were performed as previously mentioned (Raajendiran et al., 2019) to depict the hierarchical clustering of proteins and separation of adipocyte clusters based on their anatomical depot of origin.. Differentially expressed (p<0.05) proteins between the adipocyte groups (ASAT vs. VAT, GFAT vs. VAT, ASAT vs. GFAT) were then subjected to Ingenuity Pathway Analysis (IPA) (Qiagen) to determine the depot-specific differential regulation of canonical pathways and functional enrichment analyses.

## Results

### Proteome variation in adipocytes derived from human VAT, GFAT and ASAT

An overview of sample preparation and mass spectrometry analysis is presented in Figure 1. Mature adipocytes were freshly isolated from patient-matched VAT, ASAT and GFAT by collagenase digestion (Figure 1A). The patient characteristics are provided in Table S1. Total cellular proteins were extracted (Figure 1B), and a spectral library of pooled VAT, ASAT and GFAT adipocyte proteomes was generated (Figure 1C). This resulted in the identification of 4,220 proteins (FDR < 1%) of which, 3,860 proteins had peptides identified with peptide confidence ≥ 99% (Figure 2A). Individual samples from VAT, ASAT and GFAT adipocytes were then analyzed by SWATH-MS (Figure 1D), and data extracted from the spectral library, which (Figure 1E) resulted in the quantification of 2,329 proteins (peptide confidence ≥ 99%, extraction FDR < 1%) across all 15 samples (Figure 1E and 2A, Table S2). To our knowledge, this constitutes the most comprehensive proteome analysis of human adipocytes.

**Figure 2.**
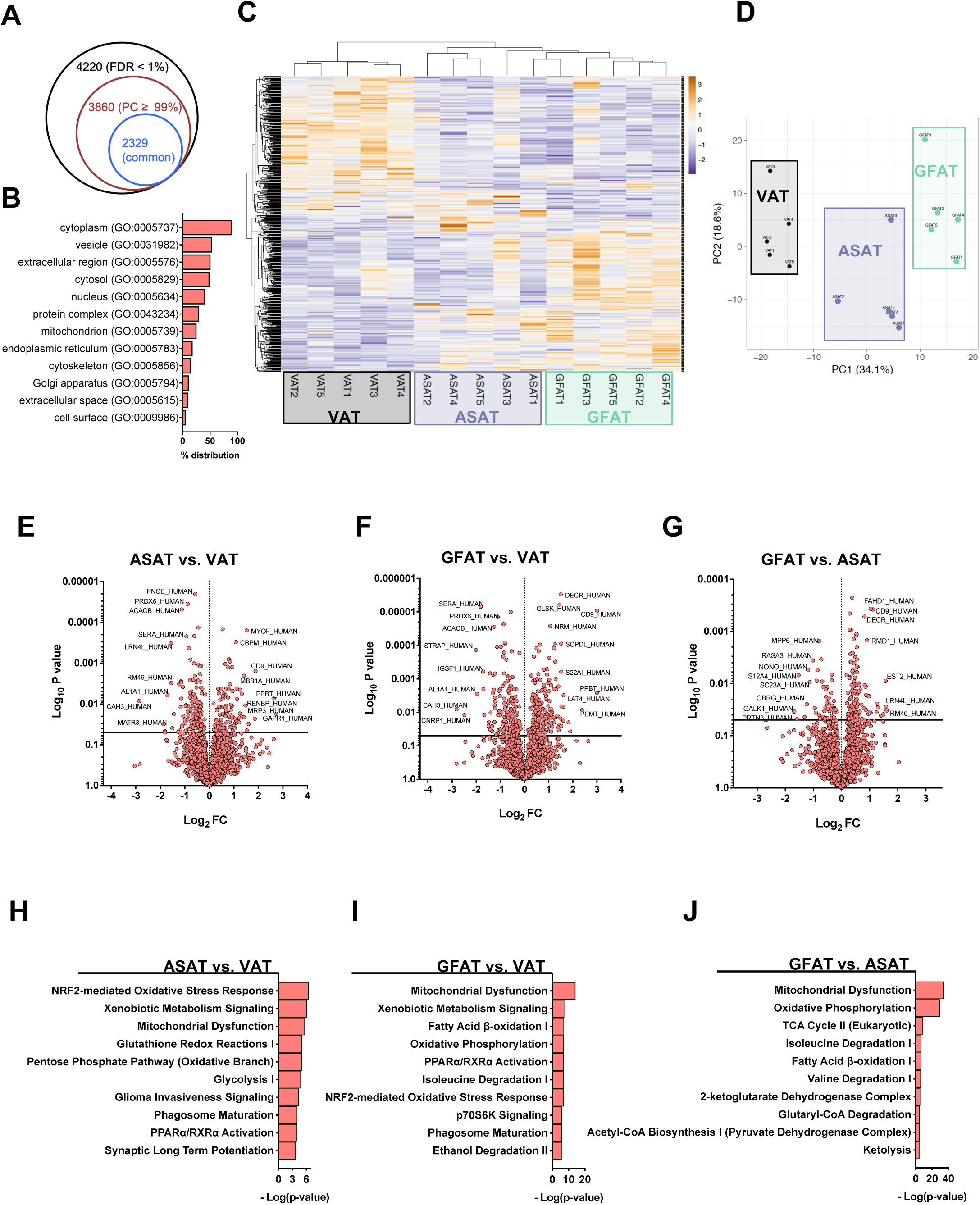
Identification and characterization of differentially expressed adipocyte proteins. **A.** Mass-spectrometry of the fractionated and non-fractionated spectral library samples resulted in the identification of 4,220 (FD < 1%) adipocyte proteins and of which 3,860 proteins had a peptide confidence of ≥ 99%. Comparing the spectra of individual samples with the spectral library resulted in the identification of 2,329 proteins (peptide confidence ≥ 99%, extraction FDR < 1%) expressed in all the test samples. **B.** Subcellular localization of the identified proteins. **C.** Heatmap and hierarchical clustering of the differentially expressed proteins in the regional adipocytes identified that VAT adipocytes are distinct from SAT (ASAT and GFAT) adipocytes. Numbers after letters denote patient numbers. **D.** Principal component analysis highlighting the clustering of regional adipocytes based on their depot of origin. Numbers after letters indicate patient numbers. **E-F.** Volcano plots showing differentially regulated proteins between the regional adipocytes. Their protein names highlight some of the highly regulated proteins between the regional adipocytes. **H-I.** Ingenuity Pathway Analysis (top 10 pathways) of differentially regulated proteins between the regional adipocytes. Top ten pathways are represented in the figures.

Gene Ontology (GO) analysis indicated that most proteins were localised in the cytoplasm (90.4%) and ~50% of the proteins were enriched with secretory functions (*i.e.*, extracellular region – GO CC:0005576) (Figure 2B). Hierarchical clustering of the differentially regulated adipocyte proteins (p < 0.05) resulted in the separation of samples into their respective anatomical locations (Figure 2C), with VAT being less similar to GFAT and ASAT. Principle component analysis (PCA) showed that the regional adipocyte proteomes were readily distinguished (Figure 2D). The differentially regulated proteins (p < 0.05, fold change (FC) ± 1.5) between the adipocyte groups are shown in Figure 2 (E-G). Comparing ASAT and VAT adipocyte proteomes, 68 proteins were upregulated in VAT adipocytes, and 70 proteins were upregulated in ASAT adipocytes (Figure 2E). When comparing GFAT and VAT, 91 proteins were upregulated in VAT adipocytes, and 105 proteins were upregulated in GFAT adipocytes (Figure 2F), while 24 proteins were upregulated, and 36 proteins were downregulated when comparing GFAT to ASAT adipocytes (Figure 2G).

### Highly abundant depot-specific proteins

Unique adipocyte proteins that were either upregulated or downregulated only in one depot when compared with the other depots are highlighted in Figure S1 (**A-F**). Of these unique proteins ALPL and GLIPR2 were highly upregulated (FC > 5) in ASAT adipocytes when compared with VAT adipocytes and the VAT adipocytes were highly enriched (FC > 5) with CA3 when compared with ASAT adipocytes (Table S2). The proteins PEMT, RCC1L, ALPL and CD9 were highly upregulated (FC > 5) in GFAT adipocytes when compared with VAT adipocytes and the VAT adipocytes were enriched (FC > 5) with CA3 and CDRIP1 when compared with GFAT adipocytes (Table S2). EMC7 and COL1A2 were highly upregulated (FC = 5) in ASAT adipocytes when compared with GFAT adipocytes and no proteins were highly upregulated (FC > 5) when comparing GFAT and ASAT adipocytes (Table S2). The functional role of some of the highly expressed proteins such as ALPL (Esteve et al., 2015), CD9 (Marcelin et al., 2017) and CA3 (Min et al., 2019) are previously reported in human adipocyte progenitor cells. The functional importance of all the highly abundant proteins (mentioned above) remain mostly unknown in mature human adipocytes.

### Bioinformatic analysis of adipocyte proteomes identifies differential regulation of metabolic pathways

Canonical pathways regulated by the adipocyte proteins were determined by Ingenuity Pathway Analysis (IPA, Qiagen). The top ten canonical pathways overrepresented by the proteins differentially regulated between the regional adipocytes are shown in Figure 2 (**H-J**). Pathways associated with mitochondrial dysfunction, oxidative phosphorylation (OXPHOS), oxidative stress, glycolysis, fatty acid oxidation, lipogenesis and branched-chain amino acid degradation were differentially regulated in the regional adipocytes. Importantly, GFAT adipocytes had an overrepresentation of fatty acid and BCAA oxidation pathways when compared with upper-body adipocytes, VAT adipocytes had an overrepresentation of oxidative stress and mitochondrial dysfunction when compared with ASAT and GFAT adipocytes and mitochondrial dysfunction was overrepresented in ASAT adipocytes when compared with GFAT adipocytes (Figure 2 H-J).

### Mitochondrial function and OXPHOS

The primary function of mitochondria is to generate ATP by integrating metabolism of various nutrients in cells (Figure 3A) (Kusminski and Scherer, 2012). IPA showed that OXPHOS was overrepresented in GFAT adipocytes compared with VAT and ASAT adipocytes (Figure 2I-J). During OXPHOS, electrons are transferred from NADH and FADH_2_ through a complex of proteins (Complex I – IV) located in the inner mitochondrial membrane (IMM), generating an electrochemical gradient across the IMM that is sufficient to drive ATP synthase (Complex V) to phosphorylate ADP to ATP in the mitochondrial matrix (Cedikova et al., 2016) (Figure 3A). Complex I - V proteins were upregulated in GFAT adipocytes when compared with ASAT adipocytes and complex I proteins were downregulated in ASAT compared with VAT adipocytes (Figure 3B). Complex IV and V proteins were downregulated in VAT adipocytes when compared with GFAT adipocytes (Figure 3B). Detailed expression profiles of the detected individual mitochondrial complex proteins are shown in Figure S2 (A-E).

**Figure 3.**
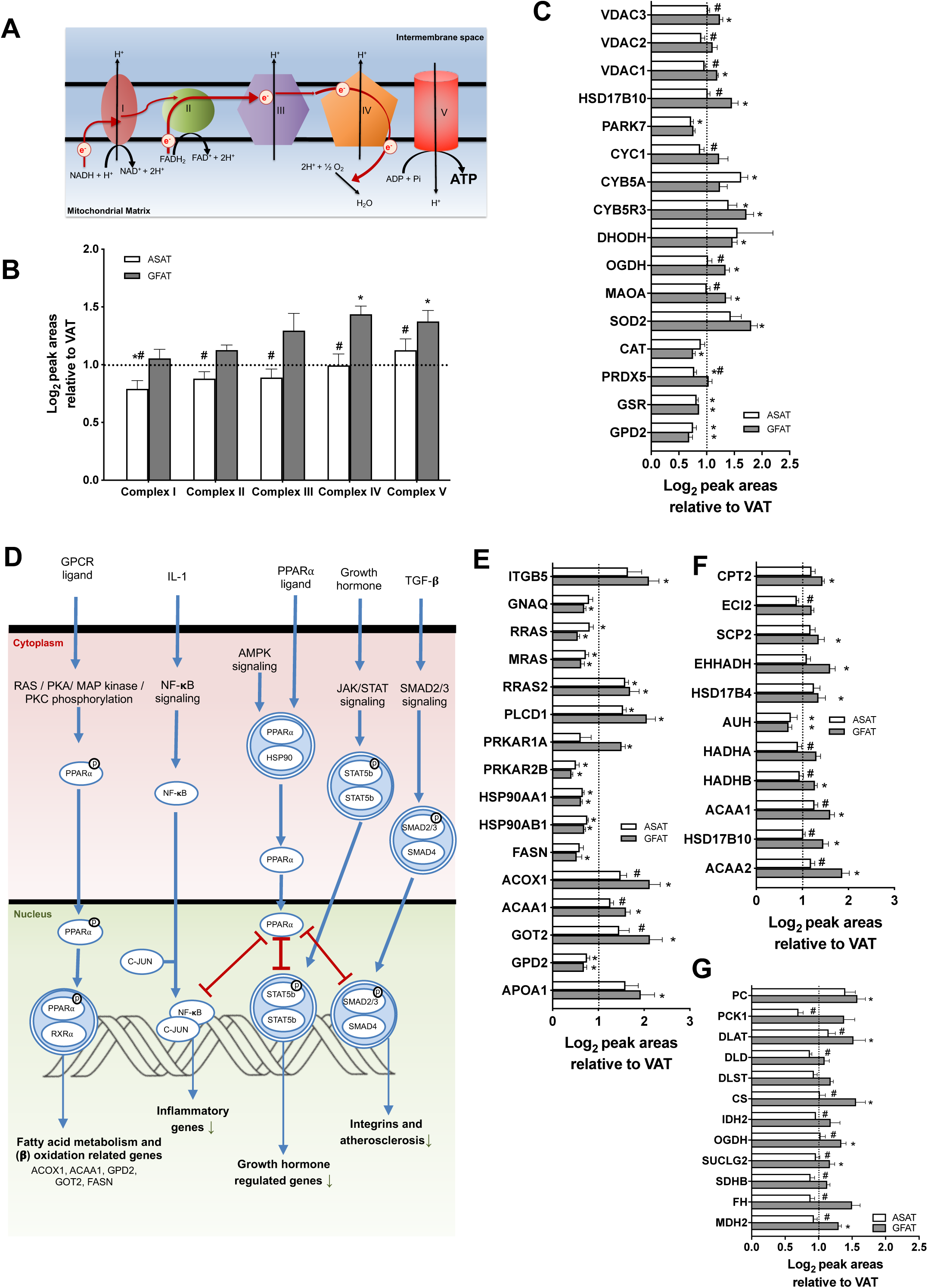
Depot-specific regulation of proteins involved in **A-B.** Oxidative phosphorylation, **C.** mitochondrial dysfunction, **D-F.** PPARα - RXRα activation, **F.** Fatty acid β-oxidation and **G.** TCA cycle pathways. Overview of **A.** oxidative phosphorylation and **D.** PPARα - RXRα activation pathways. **B-C, E-G.** Depot-specific expression levels of the proteins identified in each pathway are highlighted. Values are means ± SEM. Data normalized to VAT adipocytes for visual representation purposes. Dotted line distinguishes the up and down-regulated proteins in ASAT (open bars) and GFAT (grey bars) when compared with VAT adipocytes. # - p<0.05 ASAT vs. GFAT. * - p<0.05 ASAT/GFAT vs. VAT.

Various outer mitochondrial membrane (OMM) proteins including VDAC1, VDAC3, OGDH, HSD17B10 and MAOA were upregulated in GFAT adipocytes when compared with VAT and ASAT adipocytes (Figure 3C). Several OMM proteins such as PARK7 (vs. VAT), VDAC2 (vs. GFAT) and CYC1 (vs. GFAT) were downregulated in ASAT adipocytes (Figure 3C). GPD2, an OMM enzyme standing at the crossroads of glycolysis, OXPHOS and fatty acid metabolism (Mracek et al., 2013) was upregulated in VAT adipocytes when compared with ASAT/GFAT adipocytes, whereas CYB5R3, a major OMM redox-cycler was upregulated in SAT adipocytes when compared with VAT adipocytes (Figure 3C). Taken together, these data show that proteins regulating OXPHOS are increased in GFAT adipocytes when compared with ASAT and VAT adipocytes.

### PPARα signaling and fatty acid oxidation

PPARα is a nuclear receptor protein that responds to fasting conditions and facilitates the mobilization of endogenous fatty acids towards mitochondrial oxidation by transcribing genes involved in fatty acid metabolism, including oxidation of fatty acids to produce acetyl-CoA (Li et al., 2005) (Figure 3D). IPA identified enrichment of PPARα - RXRα activation in SAT adipocytes when compared with VAT adipocytes (Figure 2H-I). Proteins of the upstream signaling pathways and activators of PPARα such as PRKAR1A (enhanced cAMP affinity, PKA signaling), RRAS2 (RAS signaling), PLCD1 and GNAQ (MAPK / PKC signaling) were enriched in SAT compared with VAT adipocytes. In contrast, the VAT adipocytes were enriched with the nuclear receptor repressor proteins HSP90AA1 and HSP90AB1, PRKAR2B (decreased cAMP affinity, PKA signaling) and cAMP degrading proteins such as YWHAQ, YWHAG and YWHAE when compared with SAT adipocytes (Figure 3E, Table S1). Consistent with potential activation of PPARα in SAT adipocytes, downstream fatty acid oxidation related proteins such as ACOX1, ACAA, GOT2 and APOA1 (vs. VAT) were increased only in GFAT adipocytes when compared with upper-body adipocytes (Figure 3D). IPA also indicated that the fatty acid β-oxidation pathway was overrepresented in GFAT adipocytes (Figure 2I-J) due to enhanced expression of proteins including ACAA1, ACAA2, HSD17B10, HSD17B4 (vs. VAT), EHHADH (vs. VAT and p=0.06 vs. ASAT), SCP2 (vs. VAT), HADHB, HADHB (vs. ASAT, p=0.08 vs. VAT) and ECI2 (vs. ASAT and p=0.06 vs. VAT) when compared with upper body adipocytes (Figure 3F). CPT2, which is involved in mitochondrial fatty acid transport, was also upregulated in GFAT adipocytes when compared with VAT adipocytes (Figure 3F). Together, the data suggest that GFAT adipocytes have an enhanced capacity for PPARα activity and fatty acid oxidation when compared with the upper-body ASAT and VAT adipocytes.

### TCA cycle

Acetyl CoA derived from fatty acid oxidation and glycolysis is used by the tri-carboxylic acid cycle (TCA) for energy production. In line with the increased expression of glycolytic proteins, expression of enzymes of the PDH complex including DLAT (vs. VAT), DLD (vs. ASAT) and DBT (vs. ASAT) were upregulated in GFAT adipocytes (Figure 4E). In addition, a rate-limiting enzyme of the TCA cycle, citrate synthase (CS), and other TCA cycle enzymes including SDHB (vs. ASAT), FH (vs. ASAT), OGDH, DLST and MDH2 were upregulated in GFAT adipocytes when compared with upper-body VAT and ASAT adipocytes (Figure 4E). GFAT adipocytes were also enriched with the mitochondrial enzyme pyruvate carboxylase (PC), which is responsible for the irreversible conversion of pyruvate into oxaloacetate to replenish the TCA cycle (Figure 4E). The metabolome data showed no difference in the amount of pyruvate (Figure 5A), but some of the TCA cycle intermediates such as malate, fumarate and succinate were increased in VAT adipocytes. Whether this indicates reduced TCA flux when compared with GFAT adipocytes is unclear (Figure 5B). The results here suggest that the TCA cycle capacity is increased in GFAT adipocytes when compared with upper-body adipocytes.

**Figure 4.**
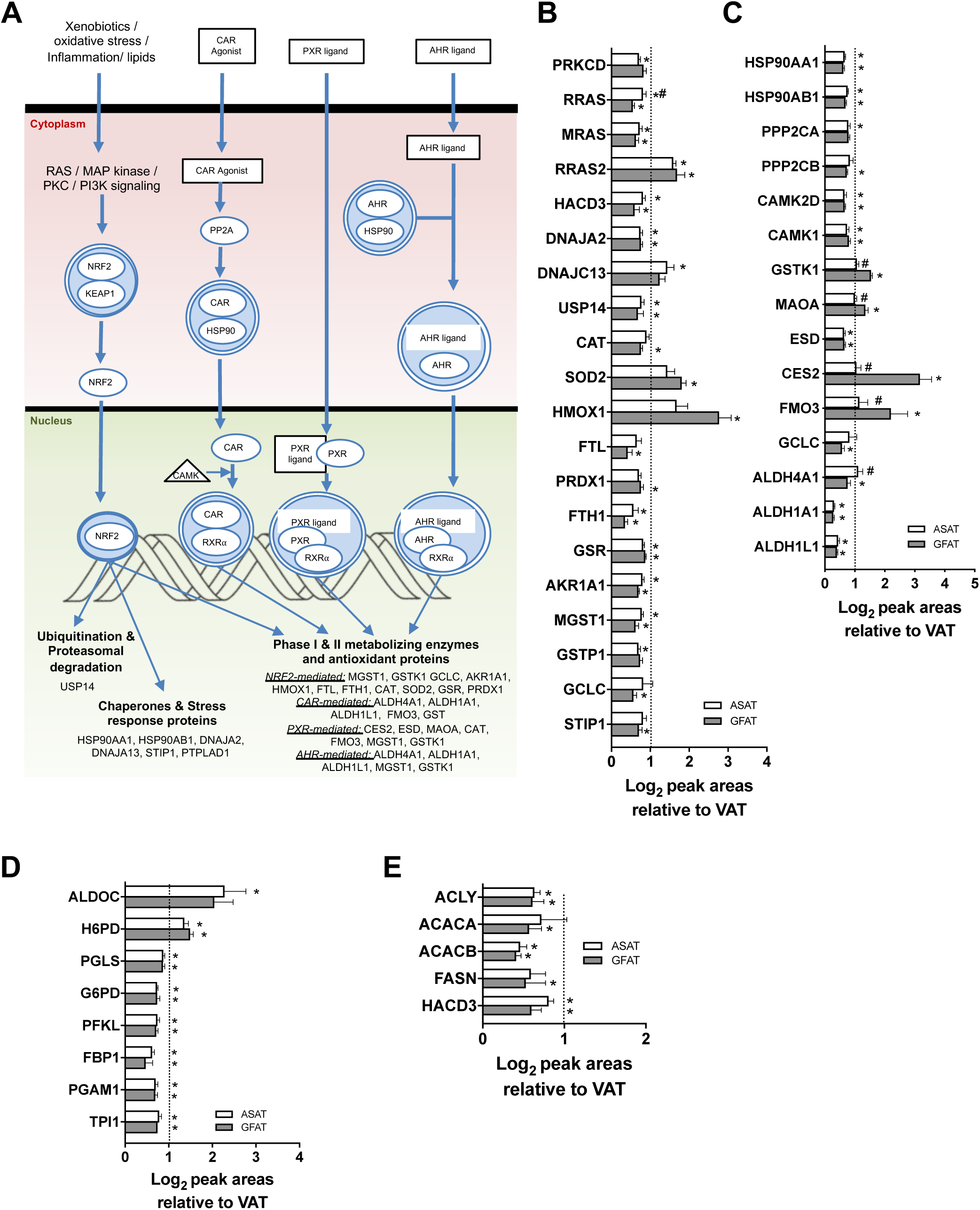
**A.** Overview and depot-specific expression levels of the proteins identified in **B.** NRF-2 mediated oxidative stress and **B-C.** Xenobiotic metabolism pathways. Values are means ± SEM. Data normalized to VAT adipocytes for visual representation purposes. Dotted line distinguishes the up and down-regulated proteins in ASAT (open bars) and GFAT (grey bars) when compared with VAT adipocytes. # - p<0.05 ASAT vs. GFAT. * - p<0.05 ASAT/GFAT vs. VAT.

**Figure 5.**
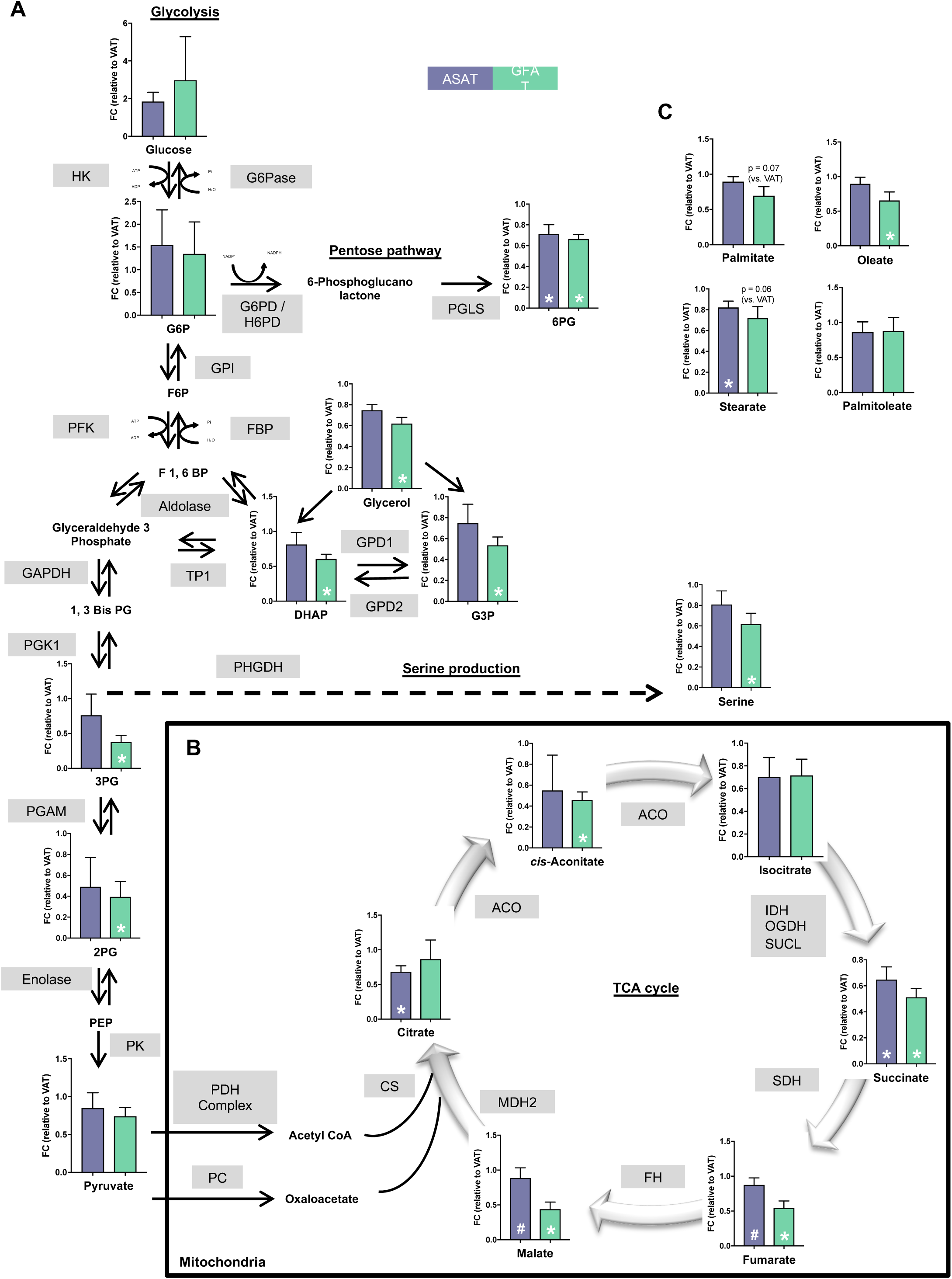
Overview and abundance of detected metabolites of **A.** glycolysis and the associated branches, including pentose and serine production and **B.** TCA cycle pathways. **C.** Metabolome data of fatty acids identified in the regional adipocytes. The metabolites and the enzymes of the pathway are abbreviated. Data normalized to VAT adipocytes and values are means ± SEM. # - p<0.05 ASAT vs. GFAT. * - p<0.05 ASAT/GFAT vs. VAT.

### NRF-2 mediated oxidative stress response and xenobiotic metabolism signaling

An imbalance between the production and scavenging of reactive oxygen species (ROS) leads to cellular oxidative stress. Pathway analysis predicted overrepresentation of the NRF2 mediated oxidative stress pathway in VAT adipocytes when compared with SAT adipocytes (Figure 2H-I). NRF2 is a transcription factor required for the induction of several antioxidant proteins (Itoh et al., 1999). The upstream signaling proteins that phosphorylate and facilitate the NRF2 nuclear translocation such as MRAS, RRAS and PRKCD were enriched in VAT adipocytes when compared with SAT adipocytes (Figure 4B-C). The key mitochondrial antioxidant enzyme SOD2 (vs. VAT) was upregulated in GFAT adipocytes and VAT adipocytes were enriched with CAT (vs GFAT) (Figure 4A-C). Additional stress response proteins such as USP14, DNAJA2, DNAJC13 (vs. ASAT), HACD3, STIP1, HSP90AA1, HSP90AB1, the phase I/II metabolizing enzymes GCLC, MGST1, GSTP1 and the antioxidant proteins FTL, FTH1, GSR, PRDX1 and CAT were upregulated in VAT adipocytes when compared with SAT adipocytes (Figure 4A-B). Overall, many antioxidant proteins are enriched in the VAT adipocytes when compared with SAT adipocytes.

The xeno-sensing nuclear receptors such as CAR, AHR and PXR also modulate the expression of stress response proteins (Figure 4A) and the xenobiotic metabolism signaling pathway was differentially regulated between the VAT and SAT adipocytes (Figure 2H-I). The heat shock proteins HSP90AA1, HSP90AB1 and DNAJA2 sequester CAR and AHR in the cytoplasm (Denis et al., 1988; Kawamoto et al., 1999) to prevent their nuclear activity and were upregulated in VAT when compared with SAT adipocytes. However, PXR-mediated (Figure 4A) expression of phase I metabolizing enzymes such as MAOA, FMO3, CES2 and the antioxidant proteins HMOX1 (vs. VAT) and SOD2 (vs. VAT) were upregulated in GFAT adipocytes when compared with upper-body adipocytes (Figure 4A-C). The data suggest that the regional adipocytes selectively activate specific xenobiotic metabolic processes.

### Glucose metabolism and the Pentose Phosphate Pathway

Upon entry to cells, glucose is converted to pyruvate via glycolysis, which enables entry and further metabolism in the mitochondria. Of the ten glycolytic enzymes, PFKL, TP1, FBP1 and PGAM1 (Figure 4D) were upregulated in VAT adipocytes when compared with SAT adipocytes. The intermediates and derivatives of the glycolytic pathway are used for various cellular biosynthetic processes, including the pentose phosphate pathway (PPP) which shunts glucose-6-phosphate into the production of pentose phosphates. The PPP was overrepresented in VAT adipocytes when compared with SAT adipocytes with upregulation of the rate-limiting enzymes G6PD and PGLS, which convert glucose-6-phosphate into ribulose-5-phosphate (Figure 4D). Alternatively, SAT adipocytes are enriched with H6PD when compared with VAT adipocytes (Figure 4D). Consistent with these protein changes, metabolomic analysis confirmed that several glycolytic intermediates including dihydroxyacetone phosphate (DHAP), glycerol-3-phosphate (G3P), 3-phosphoglycerate (3PG), 2-phosphoglycerate (2PG) and the PPP intermediate 6-phosphogluconic acid (6PG) were increased in VAT adipocytes when compared with GFAT adipocytes (Figure 5A). Together, these proteomic and metabolomic results indicate that glycolysis and PPP are upregulated in VAT adipocytes compared with SAT adipocytes.

### De novo lipogenesis

Acetyl-CoA (mostly derived from glucose) is also directed towards *de novo* synthesis of fatty acids that are mostly stored within cellular triglyceride and phospholipids. VAT adipocytes had an increased expression of ATP-citrate lyase (ACLY), the enzyme responsible for the cytosolic synthesis of acetyl-CoA from citrate, and increased content of other lipogenic enzymes including ACACA (vs. GFAT), ACACB, FASN (vs. GFAT) and HACD3 when compared with SAT adipocytes (Figure 4E). The concentration of the direct lipogenic product palmitate tended (p = 0.07) to be increased in VAT adipocytes when compared with ASAT and GFAT adipocytes (Figure 5C), although it is noted that this fatty acid can also be derived from the circulation and triglyceride hydrolysis. Together, our results highlight that VAT adipocytes have an increased capacity for glycolysis and *de novo* lipogenesis when compared with adipocytes from ASAT and GFAT.

### Amino acid metabolism

Adipose tissue is a major site of branched-chain amino acid (BCAA) metabolism (Herman et al., 2010; Rosenthal et al., 1974; Suryawan et al., 1998). BCAA catabolism is subdivided into common upstream and different downstream pathways (Figure 6A). The common pathway involves deamination of BCAAs into α-ketoacids by BCAT2 and irreversible oxidative decarboxylation of α-ketoacids by the rate-limiting branched-chain -keto acid dehydrogenase (BKCD) complex into respective acyl-CoA derivatives. BCAT2 protein content was similar in the anatomically distinct adipocytes whereas the contents of lipoamide acyltransferase (DBT) and dihydrolipoamide dehydrogenase (DLD), components of the branched-chain -keto acid dehydrogenase complex are enriched in the proteome of GFAT adipocytes when compared with ASAT adipocytes (Figure 6B). The acyl-CoA derivatives can be routed through different catabolic processes deriving substrates such as succinyl CoA and acetyl CoA, which can feed the TCA cycle. These downstream BCAA catabolic pathways were overrepresented in GFAT adipocytes when compared with upper body adipocytes with increased contents of key regulatory proteins including HIBADH, HSD17B10, HADHA (vs. ASAT), HADHB, ACAD8, ACAA1, ACAA2, ACAT1, EHHADH and MCCC2 (vs. VAT) (Figure 6B). In line with the proteome analysis, some of the intermediates of BCAA-derived ketoacid oxidation were decreased in GFAT when compared with VAT adipocytes, including β-hydroxy-β-methylbutyrate, 3-hydroxyisobutyrate and 3-hydroxybutyrate (Figure 6A). Furthermore, VAT adipocytes had increased AUH, which is responsible for the hydration of 3-methylglutaconyl-CoA (a derivative of leucine degradation) to 3-hydroxy-3-methyl-glutaryl-CoA (HMG-CoA) when compared with SAT adipocytes (Figure 6B). Overall, these results suggest that upstream common BCAA degradation pathway (vs. ASAT) and the downstream BCAA degradation pathways (vs. VAT) are overrepresented in GFAT adipocytes when compared with upper-body adipocytes. We also found that PHGDH, the first enzyme involved in *de novo* serine biosynthesis, is markedly increased in VAT adipocytes compared with ASAT and GFAT adipocytes (Figure 6B), which aligns with the increased serine concentration in VAT adipocytes (Figure 5A). In addition, ASAT and GFAT adipocytes are enriched with SHMT1, an enzyme responsible for the interconversion of serine and glycine when compared with VAT adipocytes (Figure 6B). Consistent with these findings, the metabolomic analysis shows increased L-serine in VAT adipocytes when compared with GFAT adipocytes (Figure 5A). Together, these proteomic data indicate that GFAT adipocytes have increased capacity for BCAA metabolism, whereas VAT adipocytes have an increased capacity for serine metabolism.

**Figure 6.**
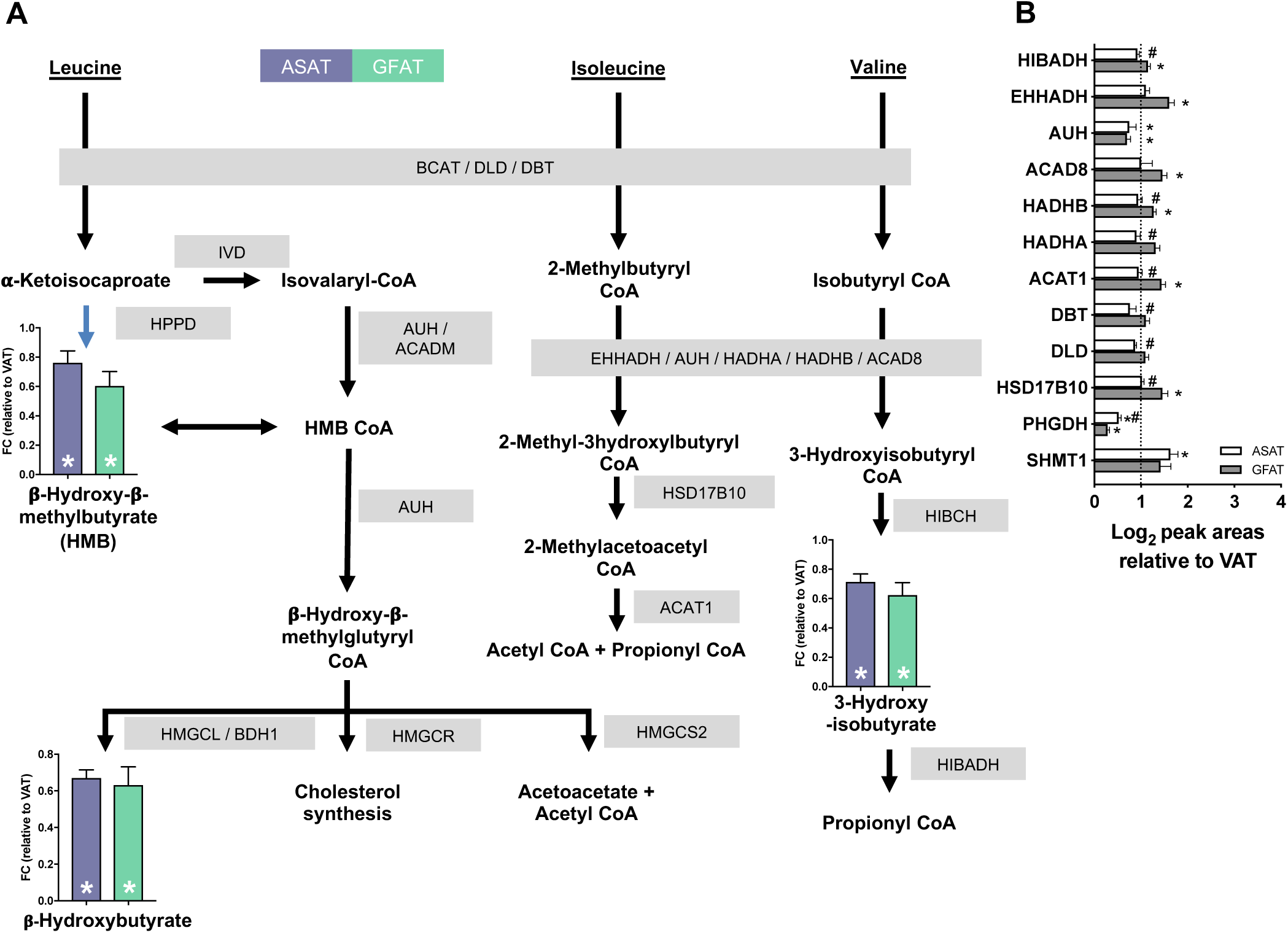
**A.** Overview and abundance of detected metabolites of branched-chain amino acid (BCAA) degradation pathway. Data normalized to VAT adipocytes. **B.** Depot-specific differential expression of BCAA degrading enzymes. **A-B.** Values are means ± SEM. **B.** Data normalized to VAT adipocytes for visual representation purposes. Dotted line distinguishes the up and down-regulated proteins in ASAT (open bars) and GFAT (grey bars) when compared with VAT adipocytes. **A-B.** # - p<0.05 ASAT vs. GFAT. * - p<0.05 ASAT/GFAT vs. VAT.

## Discussion

There remains intense interest in understanding the metabolic differences in regional adipose tissue depots and how this impacts health. This study sought to determine the proteomes of patient-matched visceral, abdominal subcutaneous and glutealfemoral adipocytes. Using recent advances in proteomics technology (Gillet et al., 2012), we identified 4,220 proteins in adipocytes, which enhanced the known proteome coverage of human adipocytes by ~1.5 fold (Xie et al., 2010). We then combined bioinformatic analysis of the adipocyte proteome with metabolomic profiling to highlight depot-specific differences in metabolic functions, that may explain, at least partially, the association of specific adipose tissue depots with increased incidence or protection from cardiometabolic disease.

Systemic lipid homeostasis is dependent on triglyceride turnover in adipocytes, which is regulated by the balance between fatty acid uptake, oxidation and esterification to triglyceride, and breakdown of triglyceride via lipolysis. Our results show that many enzymes involved with fatty acid catabolism, TCA cycle flux and oxidative phosphorylation are expressed at significantly higher levels in GFAT adipocytes compared with VAT and ASAT adipocytes. PPARα is critical for the transcription of genes that translate proteins involved in adipocyte lipolysis, fatty acid oxidation and mitochondrial biogenesis, and PPARα agonists exert weight loss (Lazennec et al., 2000; Li et al., 2005; Montgomery et al., 2019; Shalev et al., 1996). PPARα-RXRα activation was over-represented in the GFAT adipocyte proteome and the enrichment of PPARα target pathways suggest that enhanced oxidative disposal of fatty acids may be a component underpinning the so-called ‘safe’ disposal of fatty acids in GFAT. This may also explain, at least partly, why GFAT is most efficient in transporting FFAs from the circulation in women (Shadid et al., 2007). White adipose tissue plays a major role in the disposal of BCAAs (Herman et al., 2010) and their intermediates are involved in the anaplerotic replenishment of the TCA cycle, which is also important for the maintenance of fatty acid oxidation (Lerin et al., 2016). BCAA degradation pathways and associated proteins were overrepresented in GFAT adipocytes when compared with upper-body ASAT adipocytes. This is important in the context that lipid removal capacity (i.e. lipolysis accompanied by oxidation) of adipocytes is declined in obesity and dyslipidaemia (Arner et al., 2011) and that oxidative stress, mitochondrial dysfunction, and impaired BCAA degradation pathways are upregulated in ‘metabolically unhealthy’ adipocytes when compared with ‘metabolically healthy’ adipocytes (Xie et al., 2010; Xie et al., 2016). Hence, upregulation of BCAA and fatty acid disposal, as shown in GFAT adipocytes, are generally associated with a cardioprotective metabolic phenotype and is consistent with the strong association between increased lower body fat and reduced cardiovascular disease (Pinnick et al., 2012; Yim et al., 2008).

Adipose tissue has long been recognized as an important regulator of glucose metabolism, accounting for ~ 5% to 15% of whole-body glucose uptake in lean and obese individuals (Dadson et al., 2016; Virtanen et al., 2002), respectively, of which ~ 10% is used for ATP production and ~ 70% released as lactate (Marin et al., 1987). Such metabolism is consistent with an uncoupling between glycolytic flux and oxidative metabolism and or exceedingly high LDH activity in cells. Notably, proteins for oxidative phosphorylation, TCA cycle and fatty acid catabolism were uniformly decreased in VAT. Glucose uptake is increased in VAT when compared with SAT (Stolic et al., 2002) and we report upregulation of many proteins of glycolysis and the pentose phosphate pathways in VAT adipocytes. Acetyl-CoA is a product of the PDH reaction and is an important precursor for the *de novo* synthesis of the fatty acid palmitate (C16:0) (DNL) (Collins et al., 2011). DNL plays a significant role in adipose tissues, accounting for up to ~20% of the triglycerides synthesized in humans (Strawford et al., 2004). DNL is a multi-step process involving the production of malonyl CoA from acetyl CoA by the enzyme ACC, followed by condensation of acetyl CoA and malonyl CoA into palmitic acid by fatty acid synthase (FASN) in the presence of extra-mitochondrially derived NADPH (from the pentose pathway) (Goldman et al., 1963; Kornacker and Ball, 1965). Our proteomics results show that the rate-limiting pentose pathway enzyme, G6PD, the major DNL proteins ACC1 and FASN, and HACD3, an enzyme facilitating the synthesis of long-chain fatty acids, are increased in VAT adipocytes. Together, these observations support the notion that glycolytic and lipogenic capacities are increased in VAT adipocytes. This aligns with previous reports of increased glucose uptake rates in VAT vs ASAT/GFAT, particularly in obese individuals where VAT acts as an important glucose sink, and especially in insulin-resistant states where skeletal muscle glucose uptake is more impacted than adipose tissue glucose uptake (Dadson et al., 2016).

There are some important limitations to this work. We have assessed the proteome and metabolome of adipocytes obtained from fasted obese individuals and changes in the proteome during other metabolic states are unknown. Because post-translational modifications are important regulators of enzyme activity, it will be important to expand the proteomic analysis; however, the extensive tissue processing required to procure isolated adipocytes renders this technically challenging. Evidently, key changes in specific proteins and the predicted changes in cellular functions need to be formally tested using functional assays. Polar and non-polar metabolome analysis using stable isotope technology could shed interesting insights into such metabolic fluxes and the accumulation of cellular intermediates. It is also noteworthy that adipocytes were obtained from severely obese patients undergoing bariatric surgeries, and the described differences may not be conserved in lean individuals.

In conclusion, our investigation combining in-depth proteomic and metabolomic profiling of human adipocytes indicates that several highly conserved metabolic processes are likely to be differentially regulated depending on the proteome of adipocytes in various anatomical location. Increased PPARα signaling, TCA cycle flux and oxidative phosphorylation may contribute to more efficient fatty acid and BCAA catabolism in GFAT adipocytes, which may make them better equipped to counteract metabolic dysfunction. By contrast, VAT was characterized by upregulation of proteins that regulate glycolysis and lipogenesis, oxidative stress and response to stress. Finally, these data provide an important resource for understanding adipocyte functions and provides the impetus for future work examining metabolism and other cellular processes in anatomically distinct adipocytes.

## Acknowledgments

This work was supported by the National Health and Medical Research Council of Australia (NHMRC) (APP1098972). A.R. was supported by a scholarship from the Department of Physiology, Monash University. M.J.W. was supported by a Senior Research Fellowship from the NHMRC (APP1077703). R.A.T. was supported by the Victorian Cancer Agency (MCRF15023). Aspects of this research were supported by access to the Australian Proteome Analysis Facility funded by the Australian Government’s National Collaborative Research Infrastructure Scheme.

## Supplementary Figure Legends

**Figure S1.**
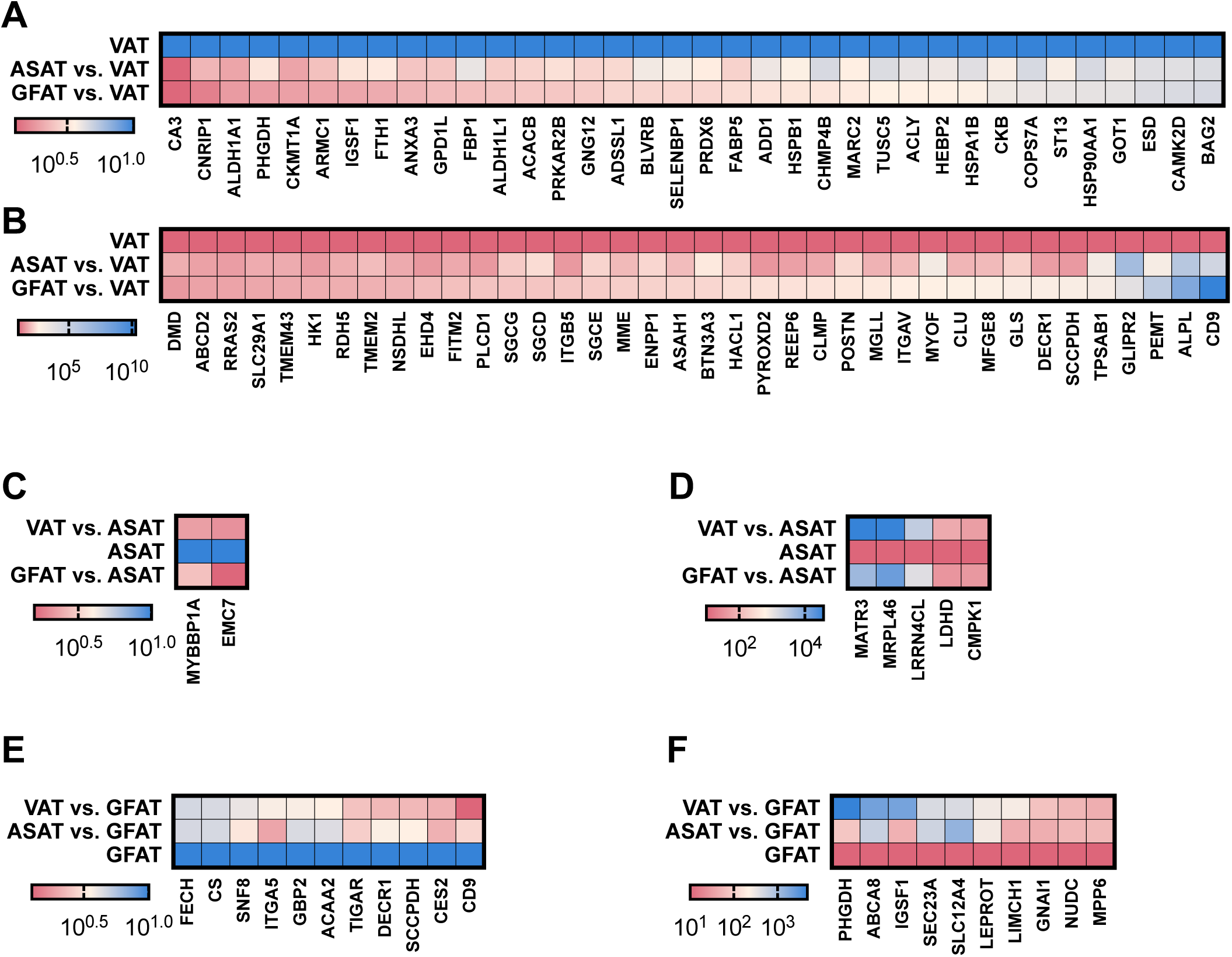
Heatmap of depot-specific unique protein. **A.** Upregulated and **B.** Downregulated proteins in VAT adipocytes vs. ASAT and GFAT adipocytes. **C.** Upregulated and **D.** Downregulated proteins in ASAT adipocytes vs. VAT and GFAT adipocytes. **E.** Upregulated and **F.** Downregulated proteins in GFAT adipocytes vs. VAT and ASAT adipocytes. Data normalized to **A-B.** VAT adipocytes, **C-D.** ASAT adipocytes and **E-F.** GFAT adipocytes for visual representation purposes.

**Figure S2.**
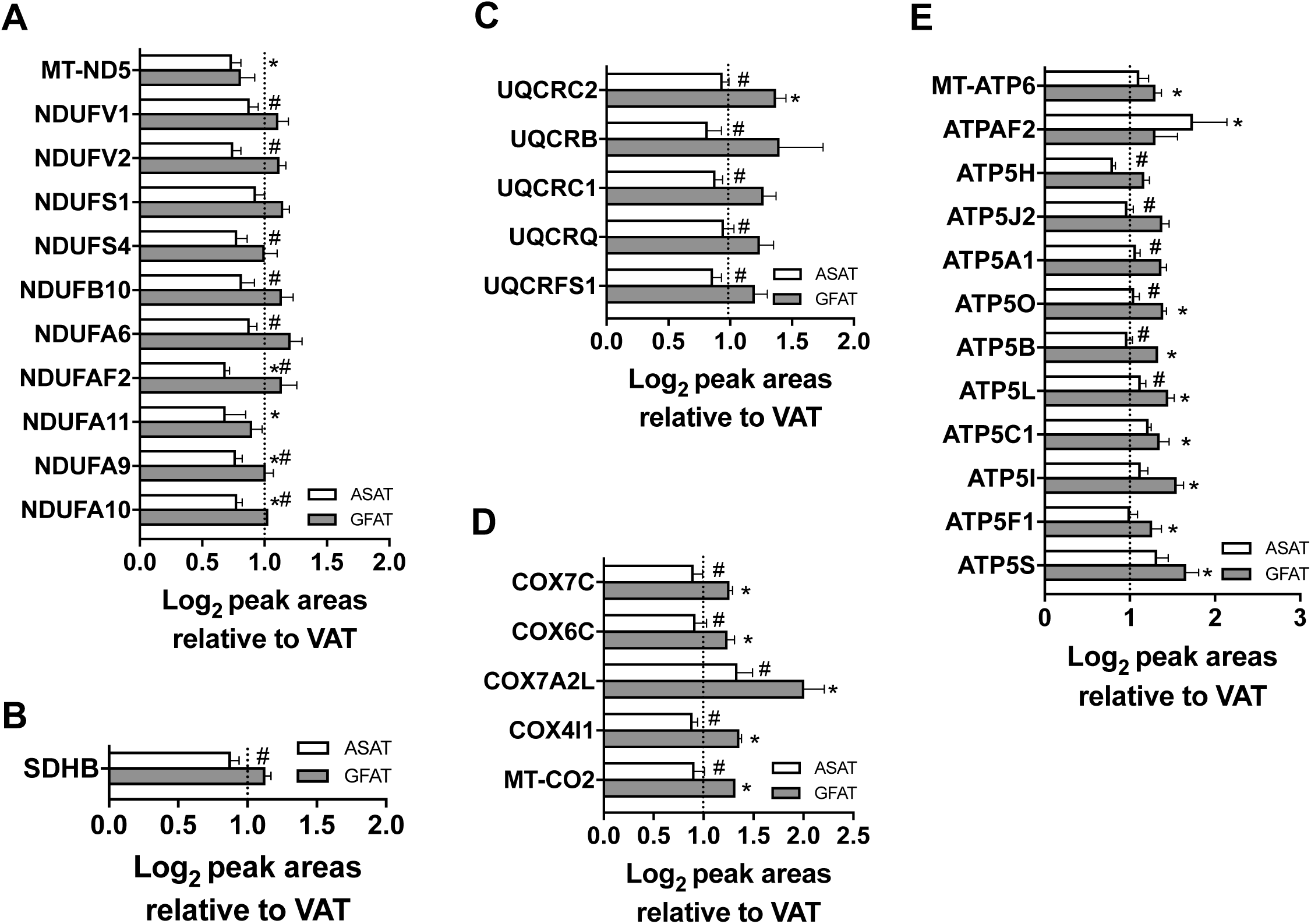
Depot-specific differential expression of mitochondrial complex proteins. **A.** Complex I, **B.** Complex II, **C.** Complex III, **D.** Complex IV and **E.** Complex V. Values are means ± SEM. Data normalized to VAT adipocytes for visual representation purposes. Dotted line distinguishes the up and down-regulated proteins in ASAT (open bars) and GFAT (grey bars) when compared with VAT adipocytes. # - p<0.05 ASAT vs. GFAT. * - p<0.05 ASAT/GFAT vs. VAT.

**Table S1.** Patient characteristics.

| Parameter |  | Proteome analysis |  | Metabolome analysis |  |
| --- | --- | --- | --- | --- | --- |
| | | Mean ± SEM | *n* | Mean ± SEM | *n* |
| Sex |  | F | 5 | F+M | 4F, 1M |
| Age | | 49.0 ± 3.4 | 5 | 48.0 ± 5.9 | 5 |
| Weight (kg) | | 135.2 ± 11.2 | 5 | 115.4 ± 1.2 | 5 |
| Height (m) | | 1.612 ± 0.02 | 5 | 1.7 ± 0.01 | 5 |
| BMI | | 52.5 ± 5.6 | 5 | 41.8 ± 0.1 | 5 |
| Insulin mU/L | | 25.0 ± 5.0 | 2 | 17.9 ± 8.3 | 5 |
| HbA1c mmol/mol | | 52.5 ± 22.5 | 2 | 36.8 ± 2.1 | 5 |
| Fasting glucose mmol/L | | 6.3 ± 1.2 | 3 | 5.3 ± 0.3 | 5 |
| c-peptide pmol/ml | | 1.2 ± 0.6 | 2 | 1.0 ± 0.2 | 4 |
| Cholesterol mmol/L | Total | 5.1 ± 0.5 | 3 | 5.2 ± 0.7 | 5 |
| | HDL | 1.4 ± 0.3 | 3 | 1.4 ± 0.2 | 5 |
| | LDL | 3.0 ± 0.4 | 3 | 2.8 ± 0.3 | 5 |
| Triglycerides mmol/L | | 1.5 ± 0.2 | 3 | 2.3 ± 0.5 | 5 |
| Total protein g/L | | 72.7 ± 1.3 | 3 | 69.3 ± 2.4 | 5 |
| Platelets (x 10<sup>9</sup>/L) | | 266.0 ± 26.5 | 3 | 298.0 ± 39.9 | 5 |
| TSH mU/L | | 2.2 ± 0.6 | 3 | 1.0 ± 0.3 | 5 |

BMI: body mass index, FPG: fasting plasma glucose, HbA1c: glycosylated hemoglobin A1c, HDL: high-density cholesterol, LDL: low-density lipoprotein, TSH: thyroid stimulating hormone.

**Table S2.** Proteins identified in the patient-matched regional adipocytes. Refer to the corresponding Excel file.

